# Recovery of Verbal Working Memory Depends on Left Hemisphere White Matter Tracts

**DOI:** 10.1101/2024.08.16.608246

**Authors:** Randi C. Martin, Junhua Ding, Ali I. Alwani, Steve H. Fung, Tatiana T. Schnur

## Abstract

Researchers propose that the recovery of language function following stroke depends on the recruitment of perilesional regions in the left hemisphere and/or homologous regions in the right hemisphere (Kiran, 2012). Many investigations of recovery focus on changes in gray matter regions (e.g., Turkeltaub et al., 2011), whereas relatively few examine white matter tracts (e.g., Schlaug et al., 2009) and none address the role of these tracts in the recovery of verbal working memory (WM). The present study addressed these gaps, examining the role of left vs. right hemisphere tracts in the longitudinal recovery of phonological and semantic WM. For 24 individuals with left hemisphere stroke, we assessed WM performance within one week of stroke (acute timepoint) and at more than six months after stroke (chronic timepoint). To address whether recovery depends on the recruitment of left or right hemisphere tracts, we assessed whether changes in WM were related to the integrity of five white matter tracts in the left hemisphere which had been implicated previously in verbal WM and their right hemisphere analogues. Behavioral results showed significant improvement in semantic but not phonological WM from the acute to chronic timepoints. Improvements in semantic WM significantly correlated with tract integrity as measured by functional anisotropy in the left direct segment of the arcuate fasciculus, inferior fronto-occipital fasciculus and inferior longitudinal fasciculus. The results confirm the role of white matter tracts in language recovery and support the involvement of the left rather than right hemisphere in the recovery of semantic WM.

## Introduction

The neural basis of the recovery of language function following brain damage is of theoretical and clinical significance. Studies on the topic have typically focused on the role of gray matter regions (Hartwigsen & Saur, 2019; Kiran, 2012), with findings indicating that perilesional left hemisphere gray matter regions and/or homologous regions in the right hemisphere support language recovery (Crinion & Price, 2005; Hartwigsen & Saur, 2019; Stefaniak et al., 2022; Turkeltaub et al., 2011; cf. Wilson & Schneck, 2021). Given the importance of the interaction among gray matter regions in cognition and language processing (Kiran & Thompson, 2019; Ribeiro et al., 2024), one would also expect that the integrity of white matter tracts connecting gray matter regions would influence recovery (Han et al., 2024; Siegel et al., 2022; Tilton-Bolowsky et al., 2024). A smaller body of research on the role of white matter tracts has again provided evidence supporting the involvement of the left and/or right hemispheres, with the integrity of tracts in one or both hemispheres relating to recovery for both spontaneous and treatment-driven language recovery (e.g., Breier et al., 2011; Schlaug et al., 2009; Sihvonen et al., 2023). These studies have most often examined the relation of white matter tracts to recovery of specific language processes such as word production (Hillis et al., 2018; McKinnon et al., 2018), sentence production (Ding & Schnur, 2022) or sentence comprehension (Xing et al., 2017). The present research addressed a novel topic – specifically, examining the role of white matter tracts in the recovery of verbal working memory (WM), which has been shown to play an important role in language comprehension and production (Martin, 2021; Martin & Schnur, 2019). In the current study, WM was assessed within the domain-specific model, which postulates separate phonological and semantic components, which play different roles in language processing (Martin, 2021; Purcell et al., 2021). Thus, the study assessed whether different white matter tracts supported the recovery of the phonological and semantic components of WM.

### Domain-specific WM

Working memory is a system that allows for the maintenance and manipulation of information over short time periods in the service of carrying out complex cognitive tasks, such as reasoning and language processing (Cowan, 2011). Considerable evidence supports the claim that there are specialized WM capacities for different domains – for instance, with separate capacities for maintaining verbal and visuo-spatial information (Baddeley, Hitch, & Allen., 2021) and, within the verbal domain, separate capacities for maintaining phonological and semantic information (Martin, Rapp, & Purcell, 2021). The current study focuses on this latter distinction between phonological and semantic information. In the domain-specific model proposed by Martin et al., (1999), there is a tight linkage between word processing and these phonological and semantic working memory capacities. That is, as a series of words are presented, their lexical-semantic and phonological representations are activated and stored in separate buffers, which can be independently disrupted by brain damage. Prior results implicate different roles for the two buffers in language comprehension and production, and thus recovery of each capacity could lead to different language outcomes (Martin & Schnur, 2019; Zahn et al., 2022; Zahn & Martin, 2024).

The initial evidence supporting the distinction between phonological and semantic WM came from case studies of brain damaged individuals who showed different patterns on tasks tapping the maintenance of phonological or semantic information for word or nonword lists. Individuals argued to have phonological WM deficits did not show standard phonological effects on WM, such as phonological similarity or word length effects, suggesting that they could not rely on phonological information for recall (Martin & He, 2004; Vallar & Baddeley, 1984). In contrast, they did show standard the standard pattern of better performance on word than nonword lists, which could be attributed to their ability to retain semantic information which was available in words but not nonwords (Papagno et al., 1991; Papagno & Vallar, 1992). Those argued to have semantic WM deficits did not show the standard advantage for memory of words over nonwords though they did show typical phonological effects on WM (Martin & He, 2004; Romani & Martin, 1999). Evidence from recognition probe tasks designed to tap the two capacities also supported the distinction in the nature of their WM deficits (Allen et al., 2012; Hamilton & Martin, 2007). The category probe task, emphasizing semantic WM, required participants to listen to word lists and judge whether a probe word was in the same semantic category (e.g., list: tulip, desk, hat; probe: rose). The rhyme probe task, emphasizing phonological WM was similar, but required participants to judge whether a probe word rhymed with any list word (e.g., list: bread table fork; probe: sled). Those with semantic WM deficits performed better on the rhyme than category probe task whereas those with phonological WM deficits showed the reverse (e.g. Martin et al., 1994). In terms of language processing, those with semantic WM deficits performed more poorly than those with phonological WM deficit on comprehension requiring integrating elements across some distance whereas those with phonological WM deficits performed more poorly than those with semantic WM deficits on verbatim sentence repetition (Martin et al., 1994; Martin & He, 2004).

More recent work taking large sample case series approaches with brain damaged individuals or individual differences approaches with healthy younger and older participants has provided factor analytic evidence of separate phonological and semantic WM factors (Zahn & Martin, 2024) and separate influences of phonological and semantic WM in language production (Martin & Schnur, 2019; Zahn et al., 2022) and comprehension (Horne et al., 2022; Tan et al., 2017; Tan & Martin, 2018; Lu, Fischer-Baum, & Martin, 2024). In general, there is evidence a greater contribution of semantic WM in both (see Martin, 2021 for review).

### Neural correlates of domain-specific WM

#### Relations to gray matter regions

In line with distinctive behavioral patterns, different gray matter regions have been identified as supporting phonological and semantic WM. Early case study findings indicated maximum lesion overlap in the left supramarginal gyrus (SMG) for those individuals showing a phonological WM deficit (see Vallar & Papagno, 2002). The role of the SMG in phonological WM has been corroborated by recent findings using multivariate lesion symptom mapping for brain damaged participants (Martin et al., 2021; Pisoni et al., 2019) and multivariate decoding methods in functional MRI studies of healthy individuals (Yue et al., 2019; Yue & Martin, 2021). It should be noted that the SMG is thought to be involved in the storage of phonological representations whereas frontal regions involved in motor planning and execution have been implicated in articulatory rehearsal of these representations (Martin et al., 2021; Vallar & Papgno, 2002; Yue et al., 2019). With respect to semantic WM, fewer studies have been carried out. However, evidence from lesion symptom mapping (Martin et al., 2021) and univariate and multivariate fMRI methods in unimpaired populations (Crosson et al., 1999; Hamilton, Martin, & Burton, 2009; Shivde & Thompson-Schill, 2004; Yue & Martin, 2021) implicates the left inferior frontal gyrus and the angular gyrus, regions which are distinct from those involved in phonological WM.

#### Relations to white matter tract integrity

With respect to the white matter correlates of verbal WM, research typically relates white matter integrity to performance on tasks such as digit, word, and nonword span (e.g. Charlton et al., 2013; Østby et al., 2011). These tasks are primarily phonological in nature, as there is little semantic information in random word and nonword lists and phonological features of the stimulus lists influence performance (e.g., phonological similarity and word length; Mueller et al., 2003). Thus, these measures are often assumed to measure the short-term retention of phonological information. Work on the white matter correlates of phonological WM typically implicates fronto-parietal tracts including the superior longitudinal fasciculus, and more specifically, the arcuate fasciculus. These findings have been replicated across many different study populations, including healthy younger and older adults (Burzynska, Preuschhof, Bäckman, Nyberg, Li, Lindenberger, & Heekeren, 2010; Charlton et al., 2013; Takeuchi et al., 2011), children (Østby et al., 2011), and people with neurological disorders (Audoin et al., 2007; Meyer et al., 2015; Sepulcre et al., 2009). In studies with healthy individuals, relations between verbal WM performance and white matter integrity have been documented for both left and right hemisphere tracts (Burzynska et al., 2010; Takeuchi et al., 2011). However, these studies often involved complex WM functions through the comparison of performance on 3-back vs. 1-back tasks (Burzynska et al., 2010) or performance on a combination of forward and backward span (Charlton et al., 2013; Østby et al., 2011; Takeuchi et al., 2011), it is possible that the involvement of the right hemisphere tracts resulted from the role of attentional and other executive processes in these WM tasks, rather than the storage of phonological and semantic information, which has been the focus of relations to language processing (Martin et al., 2021).

Recently, Horne, Ding, Martin, and Schnur (2022) examined the white matter correlates of phonological and semantic WM for 45 individuals at one year or more post-left hemisphere stroke. Performance on a semantic WM task (category probe) and phonological WM task (digit matching) was related to fractional anisotropy (FA) values for five left hemisphere white matter tracts. The tracts were selected on the grounds of prior literature implicating that tract in supporting verbal WM or because they had terminations in gray matter regions involved in phonological or semantic WM. Thus, the AF and its anterior and posterior subsegments (AAF, PAF) were included because the AAF connects the SMG, the proposed location of the phonological WM buffer (Crosson et al., 1999; Hamilton et al., 2009; Shivde & Thompson-Schill, 2004; Yue & Martin, 2021), to regions in the frontal lobe that support articulatory rehearsal (Chein & Fiez, 2001; Yue, 2018) and executive function (Smith et al., 1998) and the PAF connects the SMG to temporal lobe regions that support speech perception (Turkeltaub & Coslett, 2010). Tracts with terminations in either the inferior frontal lobe or the angular gyrus were included because of the evidence for the involvement of these regions in semantic WM. These included the direct segment of the AF (DAF), the inferior longitudinal fasciculus (ILF), inferior frontal-occipital fasciculus (IFOF), and uncinate fasciculus (UF), and the middle longitudinal fasciculus (MLF). Multiple regression analyses were carried out which used tract integrity as the dependent measure and semantic or phonological WM as the main predictor, while controlling for the other WM measure, single word comprehension, and gray matter damage at tract termini. These analyses revealed a significant independent relation of phonological WM to the integrity of the AF (overall) and the ILF. A significant independent relation to semantic WM was obtained for the PAF and the IFOF. The findings conformed to some degree to predictions, with phonological WM related to the integrity of the AF as a whole, consistent with prior findings, and semantic WM related to the integrity of the IFOF. It was suggested that the unexpected relations of the integrity of the ILF to phonological WM and the PAF to semantic WM might arise due to possible variation in terminations across individuals, with some having terminations of the ILF in the SMG and not the AG and vice versa for the PAF, given the proximity of the two regions. Thus, these findings documented the importance of left hemisphere tracts in predicting WM performance at a chronic stage post-stroke, with different tracts implicated as more important for either semantic or phonological WM. Relations to right hemisphere tract integrity were not reported, and thus this study did not address whether tracts in both hemispheres potentially contributed to performance.

### Current study

Whereas the Horne et al. study examined relations of verbal WM to white matter tract integrity at one timepoint (the chronic stage of stroke) solely in the left hemisphere, the present study examined the relation of tract integrity to longitudinal recovery as measured in changes in WM performance from an acute to a chronic timepoint within the same individuals, considering both left and right hemisphere tracts. Thus, the individuals tested in the present study were assessed on WM measures at an acute stage post-stroke (mean = 4 days, range: 2-10 days) and again at a chronic timepoint of at least 6 months post-stroke. To the extent that recovery depends solely on connections among left hemisphere brain regions, one would predict that the degree of improvement would be related only to the integrity of white matter tracts in the left hemisphere. On the other hand, if recovery depends on the functioning of homologous right hemisphere regions, one would predict that improvement would also relate to the integrity of white matter tracts in the right hemisphere – either alone or in conjunction with left hemisphere tracts. The participants in the current study were part of larger study recruiting individuals at the acute stage of left hemisphere stroke (Ding et al., 2020), which excluded those with prior strokes. Thus, these individuals did not have lesions in the right hemisphere.

We selected and dissected bilaterally two tracts for which significant relations of FA values to either phonological or semantic WM were obtained (i.e., the ILF and IFOF; Horne et al., 2022). In addition, we examined three subsegments of the arcuate fasciculus (AF) including the anterior, direct, and posterior segments following their hypothesized different contributions to language processing (Forkel et al., 2020). Thus, five tracts were examined in both the left and right hemispheres. With respect to predictions, there is, to our knowledge, no prior literature regarding white matter involvement in verbal WM recovery. One might hypothesize that those tracts uncovered in the Horne et al. study to support phonological or semantic WM would be those that would support recovery of their respective capacities, though perhaps with the involvement of right hemisphere homologues. However, several considerations might argue against these predictions. For one, if the left hemisphere tracts typically involved are very damaged, recovery using these tracts may be impossible and other left hemisphere tracts may become more involved (rather than just right hemisphere homologues; Keser, Sebastian, et al., 2020; van Hees et al., 2014). Also, it is possible that if tracts supporting, for instance, semantic WM are damaged, preserved phonological WM might be used as a backup to aid access to semantic information until gray matter or white matter changes take place to support maintenance of semantic information. Finally, although about half of the participants in the Horne et al study were recruited at the acute stage of stroke, but tested at least one year post-stroke, the other half were those recruited at the chronic stage of at least one year post-stroke and many of them were several years post-stroke. The two subsets differed substantially in lesion size, with much larger lesions for those recruited at the chronic stage (see Zahn et al., 2022 for discussion). Thus, the inclusion of those with much larger lesions in the Horne et al study may have led to a different pattern of relations of tract integrity to WM than might be observed for those with smaller lesions, like the individuals tested here. Given these considerations, it is difficult to make predictions about which specific tracts might be involved in phonological or semantic WM recovery. Because of the exploratory nature of the involvement of specific tracts, it was critical that corrections for multiple comparisons were made, which was done here using the false discovery rate (FDR) method. The results provide the first direct evidence regarding whether there is involvement of left or right hemisphere tracts, or both, in verbal WM recovery.

## Methods

### 1. Participants

As part of a larger project involving multiple comprehensive stroke centers, we consecutively recruited monolingual English speakers with left hemisphere stroke without other health conditions which impacted cognition (i.e., tumor, dementia, alcohol and/or drug dependency). Among enrolled participants, 24 (acute age = 58 years, range = 20-78 years; Education = 15 years, range = 12-23 years; 11 males; 20 right-handed) completed working memory assessments at both acute (within an average of 4 days post stroke, range: 2-12 days) and chronic (>6 months since stroke; with an average of 324 days, range: 175-480 days) timepoints as well as follow-up neuroimaging at the chronic timepoint. Left hemisphere lesions affected frontal, temporal, and parietal lobes, subcortical nuclei, white matter, and cerebellum (lesion size: average 6728 mm^3^, range 51-37085 mm^3^; Figure 1). All the participants or their caregivers provided informed consent.

**Figure 1.**
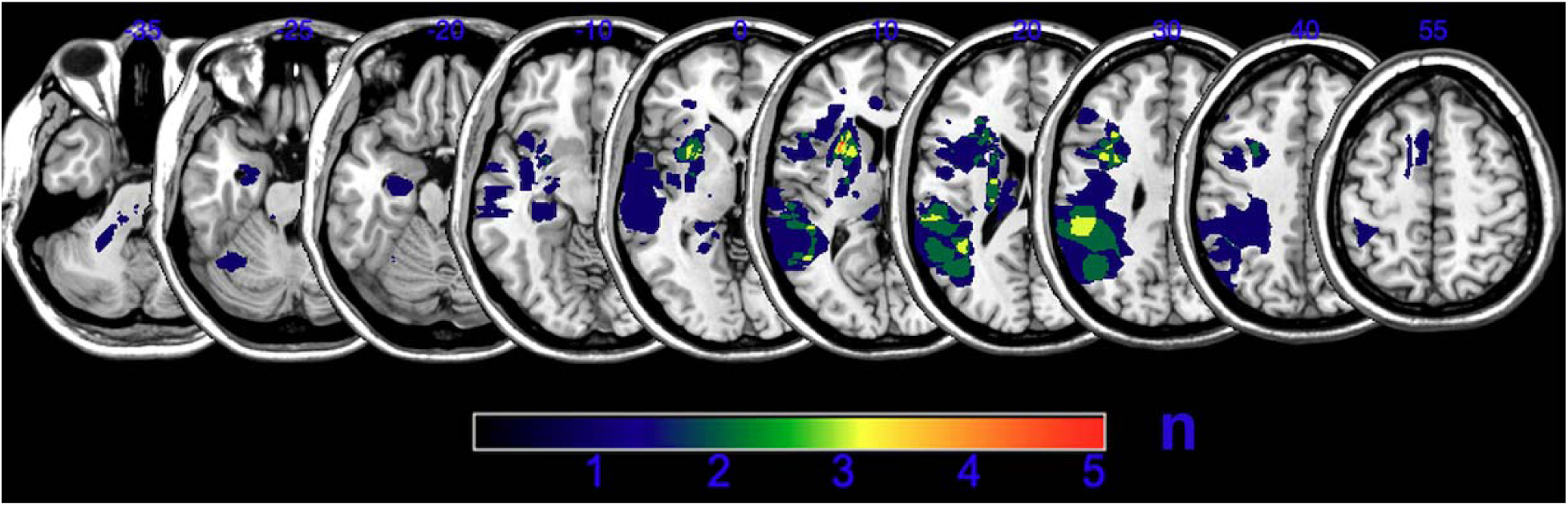
Lesion overlap. Number of individuals with lesions at each voxel.

### 2. Working memory

Digit matching span measured phonological working memory ability (Allen et al., 2012) for 22 of the 24 participants. Two participants completed the digit span task using the standard Wechsler Adult Intelligence Scale-Revised [WAIS-R] procedure (Wechsler, 1981). With respect to digit matching span, participants heard 2-digit lists (one digit/second) and then judged whether the 2 lists were the same or not. Half of the trials matched, half did not. In the ‘non-match’ trials, the second list reversed two adjacent digits and the reversed position was randomized. The list length gradually increased from 2-6 digits, with 6, 8, 6, 8 and 10 trials per list length.

A category probe task tested semantic working memory ability (Martin et al., 1994, 2021). In this task, participants heard a word list (1 word/second) and then a probe word. Participants judged whether the probe belonged to the same semantic category as any list item (e.g., matching list: tulip, desk, hat; probe: rose). Categories included animals, body parts, clothing, fruit and kitchen equipment. On half of the trials the probe matched the category of a list item and on half it did not. List length gradually increased from 1 to 4 items, with 8, 8, 12 and 16 trials per list length. For both tests of working memory, we stopped testing if accuracy fell below 75% for a particular list length. We used linear interpolation to estimate the list length corresponding to 75% accuracy.

### 3. Neuroimaging data

Due to an extended participant recruitment period, two different sets of high-resolution T1 and DTI sequences were employed at the chronic stage. Eight participants were scanned in a Philips Intera 3T scanner. The acquisition parameters were as follow: (1) diffusion tensor imaging: TR = 11098 ms, TE = 60 ms, 70 axial slices, slice thickness = 2mm, in-plane resolution: 2mm * 2mm, 32 directions plus 1 b0 volume, b-value = 800; (2) 3D T1 TFE: TR = 8.4 ms, TE = 3.9 ms, flip angle = 8 deg, 175 sagittal slices, slice thickness = 1mm, in-plane resolution: 0.94 mm * 0.94 mm. Other participants were scanned in a Siemens Prisma 3T scanner. The acquisition parameters were as follow: (1) diffusion tensor imaging: TR = 7700 ms, TE = 70 ms, 72 axial slices, slice thickness = 2mm, in-plane resolution: 2mm * 2mm, 64 directions plus 1 b0 volume, b-value = 1000; (2) 3D T1 MPRAGE: TR = 2600 ms, TE = 3.02 ms, flip angle = 8 deg, 176 sagittal slices, slice thickness = 1mm, in-plane resolution: 1 mm * 1 mm.

We used PANDA (https://www.nitrc.org/projects/panda/) (Cui et al., 2013) to perform preprocessing of diffusion weighted images. Specifically, we corrected for the eddy-current distortions, fitted the tensor model, calculated FA values, and performed deterministic tractography using the FACT algorithm, with 45°angle and 0.2 FA thresholds (Mori & van Zijl, 2002; Wang, Benner, Sorensen, & Wedeen, 2007).

Five working memory-related tracts were selected and dissected bilaterally including the anterior, direct and posterior segments of arcuate fasciculus (AF), inferior longitudinal fasciculus (ILF), and inferior frontal-occipital fasciculus (IFOF) (Horne et al., 2022; Figure 2). We used a two Region of Interest (ROI) method to isolate different tracts in TrackVis (http://trackvis.org/). The anterior segment of the AF was captured by the inferior frontal gyrus and supramarginal gyrus disk ROIs. The direct segment of the AF was captured by the inferior frontal gyrus and posterior middle temporal gyrus disk ROIs. The posterior segment of the AF was captured by the angular gyrus and posterior middle temporal gyrus disk ROIs (Catani et al., 2007). The IFOF was defined by two disk ROIs located in the ventral medial occipital lobe and anterior floor of the external/extreme capsule (Catani & Thiebaut de Schotten, 2008; Fekonja et al., 2019). The ILF was defined using the anterior temporal lobe disk ROI and the occipital lobe ROI of the IFOF (Catani & Thiebaut de Schotten, 2008; Fekonja et al., 2019). When the tract was identifiable within a participant, we extracted the FA value (Table 1).

**Figure 2.**
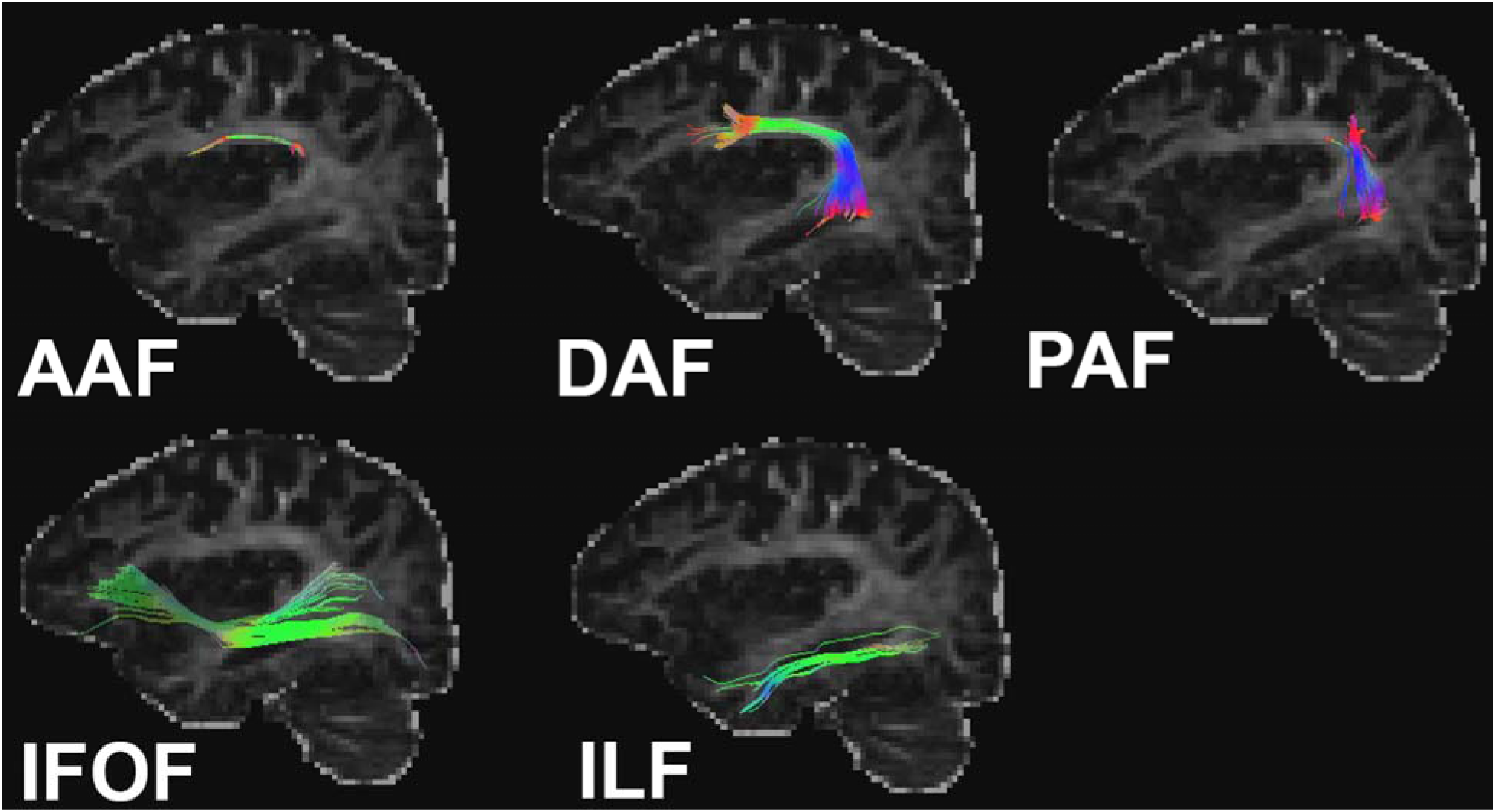
Representative tracts of interest from a single participant after left hemisphere stroke.

**Table 1.** Tract FA values.

|  | Mean | SD | Min | Max |
| --- | --- | --- | --- | --- |
| Left Hemisphere |  |  |  |  |
| AAF (n=20) | 0.39 | 0.03 | 0.33 | 0.45 |
| DAF (n=17) | 0.43 | 0.03 | 0.37 | 0.47 |
| PAF (n=18) | 0.38 | 0.03 | 0.33 | 0.42 |
| IFOF (n=24) | 0.43 | 0.03 | 0.37 | 0.47 |
| ILF (n=24) | 0.41 | 0.03 | 0.36 | 0.48 |
| Right Hemisphere |  |  |  |  |
| AAF (n=23) | 0.40 | 0.03 | 0.33 | 0.45 |
| DAF (n=9) | 0.42 | 0.03 | 0.36 | 0.46 |
| PAF (n=21) | 0.39 | 0.04 | 0.29 | 0.45 |
| IFOF (n=24) | 0.43 | 0.03 | 0.38 | 0.47 |
| ILF (n=24) | 0.41 | 0.03 | 0.34 | 0.48 |

To rule out the potential influence of gray matter damage, we calculated the gray matter damage at the tracts’ termini and used that as a control variable in our analyses. First, lesion masks were manually drawn on T1 images using ITK-snap (Yushkevich et al., 2006). Then we registered T1 images to the Colin-27 template using ANTs and normalized masks to the MNI space based on the transformation parameters (Avants et al., 2008; cf. (Ding & Schnur, 2022) for a similar approach). Using the AAL atlas to define regions, we estimated a region’s gray matter damage as the region’s intersection with the lesion masks. The AAF’s termini included the inferior frontal gyrus and supramarginal gyrus. The DAF’s termini included the inferior frontal gyrus and middle temporal gyrus. The PAF’s termini included the angular gyrus and middle temporal gyrus. The IFOF’s termini included the frontal orbital gyrus and occipital lobe. The ILF’s termini included the temporal pole and occipital lobe. We calculated total gray matter damage related to each tract as the summation of the two termini # voxels damaged divided by the summed number of total voxels included in the two termini regions.

### 4. Statistical analysis

To explore the relation between white matter tracts and working memory recovery, we conducted partial correlations between FA values of the white matter tracts and changes in WM performance from the acute to chronic timepoints. For both left and right hemisphere analyses, we controlled for acute WM baseline. In addition, for the left hemisphere, we controlled for gray matter damage of the tracts’ two termini as a result of the left hemisphere stroke. As tracts were not identified in all participants, we only analyzed tracts identified in more than half of the sample. Therefore, the right DAF was excluded from this analysis (n = 9). FDR correction was conducted within each hemisphere and within each WM task to account for multiple comparisons.

## Results

### 1. Behavioral results

Patients’ working memory performance is shown in Figure 3. Participant working memory spans and changes in span demonstrated large variability (semantic WM 2.7 ± 1.3 (acute), 3.7 ± 1.1 (chronic), 1.0 ± 1.3 (change); phonological WM 5.2 ± 1.4 (acute), 5.4 ± 1.4 (chronic), 0.2 ± 1.3 (change)). Paired t-tests indicated a significant improvement in semantic WM (t = 2.36, p = 0.03) from acute to chronic stroke, but not in phonological WM (t = 0.21, p = 0.84).

**Figure 3.**
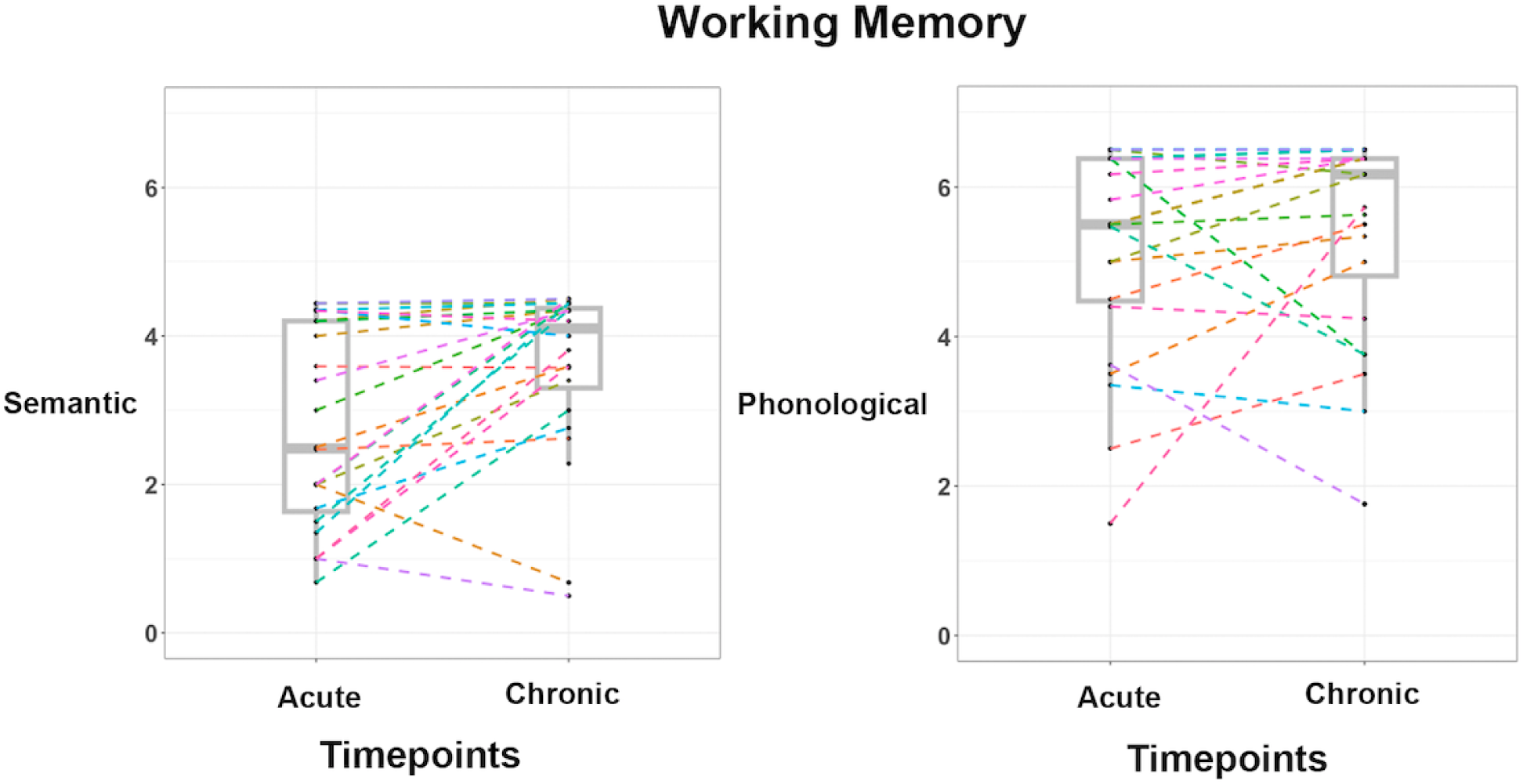
Individuals’ semantic and phonological working memory performance at acute and chronic post-stroke time points.

### 2. Relationship between FA and working memory recovery

After controlling for acute performance and gray matter LH damage of tracts’ termini, three white matter tracts’ FA significantly related with the degree of semantic WM change from acute to chronic timepoints (FDR corrected; Table 1 and Figure 4): left DAF (r = 0.70; p = 0.002), left IFOF (r = 0.52, p = 0.01) and left ILF (r = 0.56, p =0.006; no significant effects after FDR correction in the right hemisphere). Higher tract FA always led to better semantic WM recovery. No significant effects were found for phonological WM after FDR correction.

**Table 1.** Correlations between FA and WM change controlling for acute WM and tract termini damage (LH).

|  | Semantic WM |  | Phonological WM |  |
| --- | --- | --- | --- | --- |
|  | R | p | R | p |
| **LH** |  |  |  |  |
| AAF (n=20) | 0.39 | 0.11 | 0.30 | 0.23 |
| DAF (n=17) | 0.74 | **0.002** | 0.29 | 0.30 |
| PAF (n=18) | 0.33 | 0.21 | 0.52 | 0.04 |
| IFOF (n=24) | 0.51 | **0.02** | 0.35 | 0.11 |
| ILF (n=24) | 0.56 | **0.006** | 0.43 | 0.05 |
| **RH** |  |  |  |  |
| AAF (n=23) | 0.44 | 0.04 | 0.40 | 0.06 |
| PAF (n=21) | 0.30 | 0.20 | 0.35 | 0.13 |
| IFOF (n=24) | 0.44 | 0.04 | 0.26 | 0.23 |
| ILF (n=24) | 0.44 | 0.04 | 0.32 | 0.14 |

**Bold** numbers are significant after FDR correction.

**Figure 4.**
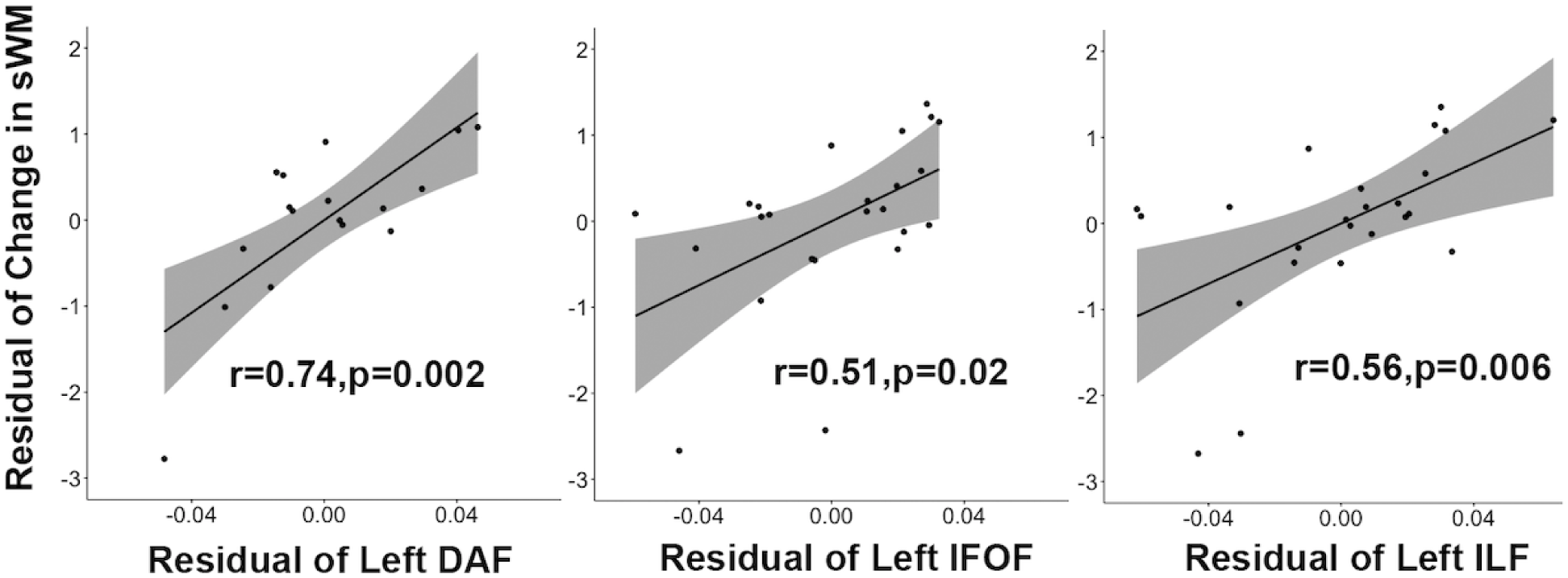
The significant effect of tracts’ FA on semantic working memory performance. sWM: semantic working memory; DAF: direct segment of arcuate fasciculus; IFOF: inferior fronto-occipital fasciculus; ILF: inferior longitudinal fasciculus.

## Discussion

While considerable research has documented changes in gray matter function in both the left and the right hemispheres during recovery of language processing following stroke (e.g., Crinion & Price, 2005; Hartwigsen & Saur, 2019; Stefaniak et al., 2022; Turkeltaub et al., 2011; cf. Wilson & Schneck, 2021), a much smaller body of research has examined the role of white matter tracts in recovery, again with some evidence indicating the involvement of both left and right hemisphere tracts (e.g. Breier et al., 2011; Schlaug et al., 2009; Sihvonen et al., 2023). Our principal aim in the current study was to determine whether both left and right hemisphere white matter tracts contribute to the recovery of phonological and semantic WM, an issue not previously addressed. For our participants with left hemisphere stroke tested at acute and chronic timepoints, semantic WM showed significant improvement, whereas phonological WM did not. Recovery of semantic WM was related to the integrity of left hemisphere tracts and not to right hemisphere tracts. For phonological WM, none of the relations to white matter tracts in either the left or right hemisphere met statistical significance. That semantic WM recovery relies exclusively on the integrity of white matter tracts in the left hemisphere contributes to current debates concerning the degree to which recruitment of the right hemisphere contributes to the recovery of function in language processes (Makin & Krakauer, 2023.; K. C. Martin et al., 2022; Wilson et al., 2023), extending the findings to verbal WM, where a tight linkage between word processing and WM capacities is assumed in the domain-specific model (Martin et al., 1999, 2021).

It should be noted that the lack of relation of phonological WM to white matter tract integrity could not be attributed to a restriction of range in phonological WM change scores. While phonological WM did not improve at the group level, there was considerable variability in the magnitude of performance change in phonological WM, with some individuals showing improvements and others showing decrements. In fact, the degree of variability in WM change was slightly higher for phonological (sd=1.29) than for semantic WM (sd=1.26). Thus, although there are many potential reasons for non-statistically significant effects, we consider a lack of variability in the change of phonological WM over time not a likely candidate.

The white matter tracts for which a significant correlation between FA values and semantic WM change scores was obtained were the left DAF, IFOF, and ILF. In the Horne et al., (2022) study which examined relations between white matter tracts and WM at a chronic timepoint, when examining pairwise correlations between FA values and semantic WM, we obtained significant correlations for all three of these tracts. However, in the multiple regression results, a significant independent contribution of semantic WM to the prediction of FA values was obtained only for the IFOF. In fact, for the ILF, a significant independent contribution was obtained for phonological WM. For the DAF, neither semantic nor phonological WM had an independent contribution. As discussed in the introduction, however, the predictions regarding the relation between tracts supporting WM at a given time point and those supporting recovery are not entirely straightforward. It is possible, for instance, that support from a tract supporting phonological WM (e.g., ILF) was helpful in providing a scaffold for the recovery of semantic WM, particularly if damage to tracts was severe – that is, providing the ability to maintain phonological representations that connect to semantic representations while mechanisms for semantic retention recover. For the DAF, it is possible that the tract supports aspects of WM common to both phonological and semantic WM (e.g., executive processes; Ribeiro et al., 2024) and the intercorrelation between the two WM measures prevented either type of WM from showing an independent contribution. Also, as noted earlier, it is possible that relations of semantic and phonological WM to tract integrity were different for the sample tested in Horne et al., which included many individuals with much larger lesions than those reported here (see Zahn et al., 2022). Only further studies having sufficient numbers of individuals at varying levels of tract disruption or lesion volume would be able to provide evidence regarding the potential contribution of these factors to tract involvement in recovery .

Another factor that may influence tract involvement during recovery is the presence or absence of language therapy. In prior studies of the role of white matter tracts in recovery, some have examined recovery without respect to the type or duration of therapy that might have been received (e.g., Keser, Meier, et al., 2020; Keser, Sebastian, et al., 2020; Schevenels et al., 2022) whereas other studies have examined the relation between behavioral treatment gains and the integrity of tracts at the outset of treatment or the relations between behavior changes and tract changes (e.g., Breier et al., 2011; Schlaug et al., 2009; Sihvonen et al., 2023). The former type often found language recovery was related to the left white matter tracts (Keser, Meier, et al., 2020). The few effects of right white matter tracts were negatively related to recovery (Keser, Sebastian, et al., 2020; c.f. Sihvonen et al., 2023). In contrast, the latter type often revealed elevated white matter changes in the right hemisphere after therapies and such changes were related to degree of behavioral recovery (Schlaug et al., 2009; Wan et al., 2014). The current study was of the former type, examining WM change without respect to treatment. Although some individuals tested here may have received treatment for language disorders between the acute and chronic testing timepoints, it is highly unlikely that this treatment was directed at WM processes. Although arguably testing and treatment for WM deficits should be incorporated into standard speech/language therapy (e.g., Greenspan et al., 2024), such is not common practice. Moreover, most studies on WM treatment have focused on phonological WM rather than both semantic and phonological WM (though see Harris et al., 2014). It is certainly possible that, if WM treatment was employed, greater involvement of right hemisphere tracts might be observed than was found here.

It should be noted, however, that a consensus seems to be emerging that there is no large-scale reorganization of language processes to the right hemisphere following brain damage (Wilson et al., 2023). Furthermore, others have argued that the terminology of “reorganization” is misleading, in that neural changes do not reflect a region taking on different functions that it did prior to brain damage (Makin & Krakauer, 2023). Instead, regions that become more involved during recovery are those that played some subsidiary role in that cognitive process prior to damage and the loss of regions more typically involved lead to greater activation in these subsidiary regions. Thus, to the extent that greater right hemisphere involvement is observed in language processes, such may reflect the enhancement of weak right hemisphere processing that persists in healthy individuals from childhood to adulthood (Martin et al., 2022). Interestingly, Makin and Krakauer suggest that greater changes in processing may occur in higher level regions that carry out more domain-general processes. On those grounds, one might have expected that WM processes would show greater right hemisphere involvement in recovery. However, as emphasized in the introduction, the type of WM that was investigated here focused on the maintenance of language representations, rather than on complex executive processes which might lead to more right hemisphere involvement.

## Limitations

Our claims regarding the absence of right hemisphere tract involvement in WM recovery are based on null results. Some of the partial correlations for right hemisphere tracts had p-values < .05, but were non-significant when controlling for multiple comparisons. Thus, it is certainly necessary to examine whether these null results would replicate in subsequent studies. Another concern is the absence of behavioral improvement in phonological WM at the group level. It is unclear why this occurred when semantic WM improved. Although mean performance was higher for the phonological WM task than the semantic WM task, the same was the case in a study using the same tasks for age-matched controls (N=13; 4.7 for category probe and 6.1 for rhyme probe) and a larger sample of acute stroke participants (N=69; 2.5 category probe and 5.0 digit matching) (Martin & Schnur, 2019). These results indicate that a higher list length is needed to equate difficulty of the two tasks. Standard deviations of mean scores and of change scores were similar for the two tasks. Thus, it does not appear that a ceiling effect prevented finding improvement on the digit matching task. Nonetheless, it would be valuable for future work to examine a number of different measures of phonological WM to determine if consistent results are found across measures.

## Conclusion

Here, we examined the degree to which white matter tracts in both left and right hemispheres contributed to the longitudinal recovery of two types of verbal working memory, phonological and semantic working memory from the acute to chronic stages of left hemisphere stroke. Tract integrity of the direct segment of the arcuate fasciculus, the inferior frontal-occipital fasciculus, and the inferior longitudinal fasciculus within the left hemisphere, but not the right hemisphere contributed to the recovery of semantic working memory. We found no significant effects associated with phonological working memory recovery in either hemisphere. To our knowledge, this is the first study relating bilateral changes in white matter tracts to the recovery of verbal working memory after acute stroke. Our study provides several additional advances over previous work. Because we assessed working memory performance within a few days after an initial left hemisphere stroke, we were able to assess the relationship between white-matter structure and function from before to after reorganization allowing a wider window for mapping the trajectory of recovery. Further, we examined this relationship within the same individuals, thus controlling for individual differences that are potential confounds in cross-sectional comparisons, and critically allowing for a more accurate assessment of causality when examining the relationship between white-matter integrity and verbal working memory. Overall, these results demonstrate that during the initial stages of recovery during the first year after stroke, the left hemisphere plays a significant role in recovery of verbal working memory, thus providing support for the hypothesis that spontaneous recovery of language-related processes depends on connections that are pre-morbidly associated with function.

## Acknowledgements

We gratefully acknowledge and thank our research subjects and their caregivers for their willingness to participate in this research. We thank Jolie Anderson, Miranda Brenneman, Cris Hamilton, Danielle Rossi, and Chia-Ming Lei for data collection. We thank Cris Hamilton for providing lesion mask demarcation. We thank the clinical neurological intensive care unit teams at the University of Texas Health Sciences Center and Memorial Hermann Hospital, The Houston Methodist Hospital, and the Baylor St. Luke’s Hospital for their assistance in patient recruitment and neurological assessment.

## References

1. Allen, C. M., Martin, R. C., & Martin, N. (2012). Relations between short-term memory deficits, semantic processing, and executive function. Aphasiology, 26(3–4), 428–461. 10.1080/02687038.2011.617436

2. Allen, R. J., Hitch, G. J., Mate, J., & Baddeley, A. D. (2012). Feature binding and attention in working memory: A resolution of previous contradictory findings. Quarterly Journal of Experimental Psychology (2006), 65(12), 2369–2383. 10.1080/17470218.2012.687384

3. Audoin, B., Guye, M., Reuter, F., Au Duong, M.-V., Confort-Gouny, S., Malikova, I., Soulier, E., Viout, P., Chérif, A. A., Cozzone, P. J., Pelletier, J., & Ranjeva, J.-P. (2007). Structure of WM bundles constituting the working memory system in early multiple sclerosis: A quantitative DTI tractography study. NeuroImage, 36(4), 1324–1330. 10.1016/j.neuroimage.2007.04.038

4. Avants, B. B., Epstein, C. L., Grossman, M., & Gee, J. C. (2008). Symmetric diffeomorphic image registration with cross-correlation: Evaluating automated labeling of elderly and neurodegenerative brain. Medical Image Analysis, 12(1), 26–41. 10.1016/j.media.2007.06.004

5. Baddeley, A., Hitch, G., & Allen, R. (2021). A Multicomponent Model of Working Memory. In R. Logie, V. Camos, & N. Cowan (Eds.), Working Memory: The state of the science (pp. 10–43). Oxford University Press. 10.1093/oso/9780198842286.003.0002

6. Breier, J. I., Juranek, J., & Papanicolaou, A. C. (2011). Changes in maps of language function and the integrity of the arcuate fasciculus after therapy for chronic aphasia. Neurocase, 17(6), 506–517. 10.1080/13554794.2010.547505

7. Burzynska, A. Z., Preuschhof, C., Bäckman, L., Nyberg, L., Li, S.-C., Lindenberger, U., & Heekeren, H. R. (2010). Age-related differences in white matter microstructure: Region-specific patterns of diffusivity. NeuroImage, 49(3), 2104–2112. 10.1016/j.neuroimage.2009.09.041

8. Catani, M., Allin, M. P. G., Husain, M., Pugliese, L., Mesulam, M. M., Murray, R. M., & Jones, D. K. (2007). Symmetries in human brain language pathways correlate with verbal recall. Proceedings of the National Academy of Sciences of the United States of America, 104(43), 17163–17168. 10.1073/pnas.0702116104

9. Catani, M., & Thiebaut de Schotten, M. (2008). A diffusion tensor imaging tractography atlas for virtual in vivo dissections. Cortex; a Journal Devoted to the Study of the Nervous System and Behavior, 44(8), 1105–1132. 10.1016/j.cortex.2008.05.004

10. Charlton, R. A., Barrick, T. R., Markus, H. S., & Morris, R. G. (2013). Verbal working and long-term episodic memory associations with white matter microstructure in normal aging investigated using tract-based spatial statistics. Psychology and Aging, 28(3), 768–777. 10.1037/a0032668

11. Chein, J. M., & Fiez, J. A. (2001). Dissociation of Verbal Working Memory System Components Using a Delayed Serial Recall Task. Cerebral Cortex, 11(11), 1003–1014. 10.1093/cercor/11.11.1003

12. Cowan, N. (2011). Working memory and attention in language use. In The handbook of psycholinguistic and cognitive processes: Perspectives in communication disorders (pp. 75–97). Psychology Press. 10.4324/9780203848005.ch4

13. Crinion, J., & Price, C. J. (2005). Right anterior superior temporal activation predicts auditory sentence comprehension following aphasic stroke. Brain: A Journal of Neurology, 128(Pt 12), 2858–2871. 10.1093/brain/awh659

14. Crosson, B., Rao, S. M., Woodley, S. J., Rosen, A. C., Bobholz, J. A., Mayer, A., Cunningham, J. M., Hammeke, T. A., Fuller, S. A., Binder, J. R., Cox, R. W., & Stein, E. A. (1999). Mapping of semantic, phonological, and orthographic verbal working memory in normal adults with functional magnetic resonance imaging. Neuropsychology, 13(2), 171–187. 10.1037/0894-4105.13.2.171

15. Cui, Z., Zhong, S., Xu, P., Gong, G., & He, Y. (2013). PANDA: A pipeline toolbox for analyzing brain diffusion images. Frontiers in Human Neuroscience, 7. 10.3389/fnhum.2013.00042

16. Ding, J., Martin, R. C., Hamilton, A. C., & Schnur, T. T. (2020). Dissociation between frontal and temporal-parietal contributions to connected speech in acute stroke. Brain: A Journal of Neurology, 143(3), 862–876. 10.1093/brain/awaa027

17. Ding, J., & Schnur, T. T. (2022). Anterior connectivity critical for recovery of connected speech after stroke. Brain Communications, 4(6), fcac266. 10.1093/braincomms/fcac266

18. Fekonja, L., Wang, Z., Bährend, I., Rosenstock, T., Rösler, J., Wallmeroth, L., Vajkoczy, P., & Picht, T. (2019). Manual for clinical language tractography. Acta Neurochirurgica, 161(6), 1125–1137. 10.1007/s00701-019-03899-0

19. Forkel, S. J., Rogalski, E., Drossinos Sancho, N., D’Anna, L., Luque Laguna, P., Sridhar, J., Dell’Acqua, F., Weintraub, S., Thompson, C., Mesulam, M.-M., & Catani, M. (2020). Anatomical evidence of an indirect pathway for word repetition. Neurology, 94(6), e594– e606. 10.1212/WNL.0000000000008746

20. Greenspan, W., Vieira, S., & Martin, N. (2024). Revealing linguistic and verbal short-term and working memory abilities in people with severe aphasia. Aphasiology, 0(0), 1–36. 10.1080/02687038.2024.2322770

21. Hamilton, A. C., & Martin, R. C. (2007). Proactive interference in a semantic short-term memory deficit: Role of semantic and phonological relatedness. Cortex; a Journal Devoted to the Study of the Nervous System and Behavior, 43(1), 112–123. 10.1016/s0010-9452(08)70449-0

22. Hamilton, A. C., Martin, R. C., & Burton, P. C. (2009). Converging functional magnetic resonance imaging evidence for a role of the left inferior frontal lobe in semantic retention during language comprehension. Cognitive Neuropsychology, 26(8), 685–704. 10.1080/02643291003665688

23. Han, Y., Jing, Y., Shi, Y., Mo, H., Wan, Y., Zhou, H., & Deng, F. (2024). The role of language-related functional brain regions and white matter tracts in network plasticity of post-stroke aphasia. Journal of Neurology, 271(6), 3095–3115. 10.1007/s00415-024-12358-5

24. Harris, L., Olson, A., & Humphreys, G. (2014). The link between STM and sentence comprehension: A neuropsychological rehabilitation study. Neuropsychological Rehabilitation, 24(5), 678–720. 10.1080/09602011.2014.892885

25. Hartwigsen, G., & Saur, D. (2019). Neuroimaging of stroke recovery from aphasia – Insights into plasticity of the human language network. NeuroImage, 190, 14–31. 10.1016/j.neuroimage.2017.11.056

26. Hillis, J. M., Ruan, A. B., Lazarus, J. E., Montgomery, M. W., & Berkowitz, A. L. (2018). Clinical Reasoning: A 48-year-old woman with confusion, personality change, and multiple enhancing brain lesions. Neurology, 90(19), e1724–e1729. 10.1212/WNL.0000000000005484

27. Horne, A., Zahn, R., Najera, O. I., & Martin, R. C. (2022). Semantic Working Memory Predicts Sentence Comprehension Performance: A Case Series Approach. Frontiers in Psychology, 13, 887586. 10.3389/fpsyg.2022.887586

28. Keser, Z., Meier, E. L., Stockbridge, M. D., & Hillis, A. E. (2020). The role of microstructural integrity of major language pathways in narrative speech in the first year after stroke. Journal of Stroke and Cerebrovascular Diseases: The Official Journal of National Stroke Association, 29(9), 105078. 10.1016/j.jstrokecerebrovasdis.2020.105078

29. Keser, Z., Sebastian, R., Hasan, K. M., & Hillis, A. E. (2020). Right Hemispheric Homologous Language Pathways Negatively Predicts Poststroke Naming Recovery. Stroke, 51(3), 1002–1005. 10.1161/STROKEAHA.119.028293

30. Kiran, S. (2012). What Is the Nature of Poststroke Language Recovery and Reorganization? ISRN Neurology, 2012, 786872. 10.5402/2012/786872

31. Kiran, S., & Thompson, C. K. (2019). Neuroplasticity of Language Networks in Aphasia: Advances, Updates, and Future Challenges. Frontiers in Neurology, 10. 10.3389/fneur.2019.00295

32. Makin, T. R., & Krakauer, J. W. (2023). Against cortical reorganisation. eLife, 12, e84716. 10.7554/eLife.84716

33. Martin, K. C., Seydell-Greenwald, A., Berl, M. M., Gaillard, W. D., Turkeltaub, P. E., & Newport, E. L. (2022). A Weak Shadow of Early Life Language Processing Persists in the Right Hemisphere of the Mature Brain. *Neurobiology of Language (Cambridge*, Mass.), 3(3), 364–385. 10.1162/nol_a_00069

34. Martin, R. C. (2021). The Critical Role of Semantic Working Memory in Language Comprehension and Production. Current Directions in Psychological Science, 30(4), 283–291. 10.1177/0963721421995178

35. Martin, R. C., Ding, J., Hamilton, A. C., & Schnur, T. T. (2021). Working Memory Capacities Neurally Dissociate: Evidence from Acute Stroke. Cerebral Cortex Communications, 2(2), tgab005. 10.1093/texcom/tgab005

36. Martin, R. C., & He, T. (2004). Semantic short-term memory and its role in sentence processing: A replication. Brain and Language, 89(1), 76–82. 10.1016/S0093-934X(03)00300-6

37. Martin, R. C., Lesch, M. F., & Bartha, M. C. (1999). Independence of Input and Output Phonology in Word Processing and Short-Term Memory. Journal of Memory and Language, 41(1), 3–29. 10.1006/jmla.1999.2637

38. Martin, R. C., Rapp, B., & Purcell, J. (2021). Domain-Specific Working Memory. In R. Logie, V. Camos, and N. Cowan (Eds.), Working Memory: The state of the science (pp. 249–281). Oxford University Press. 10.1093/oso/9780198842286.003.0009

39. Martin, R. C., & Schnur, T. T. (2019). Independent contributions of semantic and phonological working memory to spontaneous speech in acute stroke. Cortex; a Journal Devoted to the Study of the Nervous System and Behavior, 112, 58–68. 10.1016/j.cortex.2018.11.017

40. Martin, R. C., Shelton, J. R., & Yaffee, L. S. (1994). Language processing and working memory: Neuropsychological evidence for separate phonological and semantic capacities. Journal of Memory and Language, 33(1), 83–111. 10.1006/jmla.1994.1005

41. McKinnon, E. T., Fridriksson, J., Basilakos, A., Hickok, G., Hillis, A. E., Spampinato, M. V., Gleichgerrcht, E., Rorden, C., Jensen, J. H., Helpern, J. A., & Bonilha, L. (2018). Types of naming errors in chronic post-stroke aphasia are dissociated by dual stream axonal loss. Scientific Reports, 8(1), 14352. 10.1038/s41598-018-32457-4

42. Meyer, A. M., Snider, S. F., Campbell, R. E., & Friedman, R. B. (2015). Phonological Short-Term Memory in Logopenic Variant Primary Progressive Aphasia and Mild Alzheimer’s Disease. Cortex; a Journal Devoted to the Study of the Nervous System and Behavior, 71, 183–189. 10.1016/j.cortex.2015.07.003

43. Mori, S., & van Zijl, P. C. M. (2002). Fiber tracking: Principles and strategies - a technical review. NMR in Biomedicine, 15(7–8), 468–480. 10.1002/nbm.781

44. Mueller, S. T., Seymour, T. L., Kieras, D. E., & Meyer, D. E. (2003). Theoretical implications of articulatory duration, phonological similarity, and phonological complexity in verbal working memory. Journal of Experimental Psychology. Learning, Memory, and Cognition, 29(6), 1353–1380. 10.1037/0278-7393.29.6.1353

45. Østby, Y., Tamnes, C. K., Fjell, A. M., & Walhovd, K. B. (2011). Morphometry and connectivity of the fronto-parietal verbal working memory network in development. Neuropsychologia, 49(14), 3854–3862. 10.1016/j.neuropsychologia.2011.10.001

46. Papagno, C., Valentine, T., & Baddeley, A. (1991). Phonological short-term memory and foreign-language vocabulary learning. Journal of Memory and Language, 30(3), 331–347. 10.1016/0749-596X(91)90040-Q

47. Papagno, C., & Vallar, G. (1992). Phonological short-term memory and the learning of novel words: The effect of phonological similarity and item length. The Quarterly Journal of Experimental Psychology A: Human Experimental Psychology, 44A(1), 47–67. 10.1080/14640749208401283

48. Papagno, C., & Vallar, G. (1995). Verbal short-term memory and vocabulary learning in polyglots. *The Quarterly Journal of Experimental Psychology. A*, Human Experimental Psychology, 48(1), 98–107. 10.1080/14640749508401378

49. Pisoni, A., Mattavelli, G., Casarotti, A., Comi, A., Riva, M., Bello, L., & Papagno, C. (2019). The neural correlates of auditory-verbal short-term memory: A voxel-based lesion-symptom mapping study on 103 patients after glioma removal. Brain Structure & Function, 224(6), 2199–2211. 10.1007/s00429-019-01902-z

50. Purcell, J., Rapp, B., & Martin, R. C. (2021). Distinct Neural Substrates Support Phonological and Orthographic Working Memory: Implications for Theories of Working Memory. Frontiers in Neurology, 12, 681141. 10.3389/fneur.2021.681141

51. Ribeiro, M., Yordanova, Y. N., Noblet, V., Herbet, G., & Ricard, D. (2024). White matter tracts and executive functions: A review of causal and correlation evidence. Brain: A Journal of Neurology, 147(2), 352–371. 10.1093/brain/awad308

52. Romani, C., & Martin, R. (1999). A deficit in the short-term retention of lexical-semantic information: Forgetting words but remembering a story. Journal of Experimental Psychology. General, 128(1), 56–77. 10.1037//0096-3445.128.1.56

53. Schevenels, K., Gerrits, R., Lemmens, R., De Smedt, B., Zink, I., & Vandermosten, M. (2022). Early white matter connectivity and plasticity in post stroke aphasia recovery. NeuroImage. Clinical, 36, 103271. 10.1016/j.nicl.2022.103271

54. Schlaug, G., Marchina, S., & Norton, A. (2009). Evidence for Plasticity in White Matter Tracts of Chronic Aphasic Patients Undergoing Intense Intonation-based Speech Therapy. Annals of the New York Academy of Sciences, 1169, 385–394. 10.1111/j.1749-6632.2009.04587.x

55. Sepulcre, J., Masdeu, J. C., Pastor, M. A., Goñi, J., Barbosa, C., Bejarano, B., & Villoslada, P. (2009). Brain pathways of verbal working memory: A lesion–function correlation study. NeuroImage, 47(2), 773–778. 10.1016/j.neuroimage.2009.04.054

56. Shivde, G., & Thompson-Schill, S. L. (2004). Dissociating semantic and phonological maintenance using fMRI. Cognitive, Affective & Behavioral Neuroscience, 4(1), 10–19. 10.3758/cabn.4.1.10

57. Siegel, J. S., Shulman, G. L., & Corbetta, M. (2022). Mapping correlated neurological deficits after stroke to distributed brain networks. Brain Structure & Function, 227(9), 3173– 3187. 10.1007/s00429-022-02525-7

58. Sihvonen, A. J., Vadinova, V., Garden, K. L., Meinzer, M., Roxbury, T., O’Brien, K., Copland, D., McMahon, K. L., & Brownsett, S. L. E. (2023). Right hemispheric structural connectivity and poststroke language recovery. Human Brain Mapping, 44(7), 2897– 2904. 10.1002/hbm.26252

59. Smith, E. E., Jonides, J., Marshuetz, C., & Koeppe, R. A. (1998). Components of verbal working memory: Evidence from neuroimaging. Proceedings of the National Academy of Sciences of the United States of America, 95(3), 876–882. 10.1073/pnas.95.3.876

60. Stefaniak, J. D., Geranmayeh, F., & Lambon Ralph, M. A. (2022). The multidimensional nature of aphasia recovery post-stroke. Brain, 145(4), 1354–1367. 10.1093/brain/awab377

61. Takeuchi, H., Taki, Y., Sassa, Y., Hashizume, H., Sekiguchi, A., Fukushima, A., & Kawashima, R. (2011). Verbal working memory performance correlates with regional white matter structures in the frontoparietal regions. Neuropsychologia, 49(12), 3466–3473. 10.1016/j.neuropsychologia.2011.08.022

62. Tan, Y., & Martin, R. C. (2018). Verbal short-term memory capacities and executive function in semantic and syntactic interference resolution during sentence comprehension: Evidence from aphasia. Neuropsychologia, 113, 111–125. 10.1016/j.neuropsychologia.2018.03.001

63. Tan, Y., Martin, R. C., & Van Dyke, J. A. (2017). Semantic and Syntactic Interference in Sentence Comprehension: A Comparison of Working Memory Models. Frontiers in Psychology, 8, 198. 10.3389/fpsyg.2017.00198

64. Tilton-Bolowsky, V., Stockbridge, M. D., & Hillis, A. E. (2024). Remapping and Reconnecting the Language Network after Stroke. Brain Sciences, 14(5), 419. 10.3390/brainsci14050419

65. Turkeltaub, P. E., & Coslett, H. B. (2010). Localization of sublexical speech perception components. Brain and Language, 114(1), 1–15. 10.1016/j.bandl.2010.03.008

66. Turkeltaub, P. E., Messing, S., Norise, C., & Hamilton, R. H. (2011). Are networks for residual language function and recovery consistent across aphasic patients? Neurology, 76(20), 1726–1734. 10.1212/WNL.0b013e31821a44c1

67. Vallar, G., & Baddeley, A. D. (1984). Fractionation of working memory: Neuropsychological evidence for a phonological short-term store. Journal of Verbal Learning & Verbal Behavior, 23(2), 151–161. 10.1016/S0022-5371(84)90104-X

68. Vallar, G., & Papagno, C. (2002). Neuropsychological impairments of verbal short-term memory. In A. D. Baddeley, M. D. Kopelman, & B. A. Wilson (Eds.), The Handbook of Memory Disorders (pp. 249–270). John Wiley & Sons.

69. van Hees, S., McMahon, K., Angwin, A., de Zubicaray, G., Read, S., & Copland, D. A. (2014). Changes in white matter connectivity following therapy for anomia post stroke. Neurorehabilitation and Neural Repair, 28(4), 325–334. 10.1177/1545968313508654

70. Wan, C. Y., Zheng, X., Marchina, S., Norton, A., & Schlaug, G. (2014). Intensive therapy induces contralateral white matter changes in chronic stroke patients with Broca’s aphasia. Brain and Language, 136, 1–7. 10.1016/j.bandl.2014.03.011

71. Wang, R., Benner, T., Sorensen, A. G., & Wedeen, V.J. (2007). Diffusion toolkit: A software package for diffusion imaging data processing and tractography.

72. Wechsler, D. (1981). Manual for the Wechsler adult intelligence scale-revised. Psychological Corp.

73. Wilson, S. M., Entrup, J. L., Schneck, S. M., Onuscheck, C. F., Levy, D. F., Rahman, M., Willey, E., Casilio, M., Yen, M., Brito, A. C., Kam, W., Davis, L. T., de Riesthal, M., & Kirshner, H. S. (2023). Recovery from aphasia in the first year after stroke. Brain: A Journal of Neurology, 146(3), 1021–1039. 10.1093/brain/awac129

74. Wilson, S. M., & Schneck, S. M. (2021). Neuroplasticity in post-stroke aphasia: A systematic review and meta-analysis of functional imaging studies of reorganization of language processing. *Neurobiology of Language (Cambridge*, Mass.), 2(1), 22–82. 10.1162/nol_a_00025

75. Xing, S., Lacey, E. H., Skipper-Kallal, L. M., Zeng, J., & Turkeltaub, P. E. (2017). White Matter Correlates of Auditory Comprehension Outcomes in Chronic Post-Stroke Aphasia. Frontiers in Neurology, 8. 10.3389/fneur.2017.00054

76. Yue, Q. (2018). Evaluating the Buffer vs. Embedded Processes Accounts of Verbal Short-term Memory by Using Multivariate Neuroimaging and Brain Stimulation Approaches [Thesis, Rice University]. https://scholarship.rice.edu/handle/1911/107974

77. Yue, Q., & Martin, R. C. (2021). Maintaining verbal short-term memory representations in non-perceptual parietal regions. Cortex; a Journal Devoted to the Study of the Nervous System and Behavior, 138, 72–89. 10.1016/j.cortex.2021.01.020

78. Yue, Q., Martin, R. C., Hamilton, A. C., & Rose, N. S. (2019). Non-perceptual Regions in the Left Inferior Parietal Lobe Support Phonological Short-term Memory: Evidence for a Buffer Account? Cerebral Cortex (New York, N.Y.: 1991), 29(4), 1398–1413. 10.1093/cercor/bhy037

79. Yushkevich, P. A., Piven, J., Hazlett, H. C., Smith, R. G., Ho, S., Gee, J. C., & Gerig, G. (2006). User-guided 3D active contour segmentation of anatomical structures: Significantly improved efficiency and reliability. NeuroImage, 31(3), 1116–1128. 10.1016/j.neuroimage.2006.01.015

80. Zahn, R., & Martin, R. C. (2024). The role of input vs. output phonological working memory in narrative production: Evidence from case series and case study approaches. Cognitive Neuropsychology, 1–23. 10.1080/02643294.2024.2366467

81. Zahn, R., Schnur, T. T., & Martin, R. C. (2022). Contributions of semantic and phonological working memory to narrative language independent of single word production: Evidence from acute stroke. Cognitive Neuropsychology, 39(5–8), 296–324. 10.1080/02643294.2023.2186782

