## supplementary materials for "Recovery of Verbal Working Memory Depends on Left Hemisphere White Matter Tracts"

```
setwd("~/working_memory")
library(readxl)
library(dplyr)
library(reshape2)
library(ggpubr)
library(reshape2)
library(ggplot2)
library(sjPlot)
library(sjmisc)
library(lmerTest)
library(effects)
library(lme4)
library(broom.mixed)
library(gridExtra)
library(rstatix)
library(ppcor)

Beh<-read_excel('./fa_volume.xlsx',sheet = 'behavior')
Data1<-subset(Beh,subset=group %in% c(3,5,6,7))
Data1<-subset(Data1,subset = time %in% c('1yr'))
Data2<-subset(Beh,subset=group %in% c(2,4))
Data2<-subset(Data2,subset = time %in% c('6mo'))
Data3<-subset(Beh,subset= group %in% c(2,3,4,5,6,7))
Data3<-subset(Data3,subset = time %in% c('acute'))
Data<-rbind(Data1,Data2,Data3)

Fa<-read_excel('./fa_volume.xlsx',sheet = 'fa')
Fa_name<-colnames(Fa[-c(1,2,22)])

Fa[Fa==0]<-NA

Fa1<-subset(Fa,subset=group %in% c(3,5,6,7))
Fa1<-subset(Fa1,subset = time %in% c('1yr'))
Fa2<-subset(Fa,subset=group %in% c(2,4))
Fa2<-subset(Fa2,subset = time %in% c('6mo'))
Fa_all<-rbind(Fa1,Fa2)

Data_fa<-left_join(subset(Data,subset = time %in% c('acute')),
                   subset(Data,subset = time %in% c('6mo','1yr')),'sub')
Data_fa<- left_join(Data_fa,Fa_all,'sub')

Data_fa$pWM_change<-Data_fa$pWM.y - Data_fa$pWM.x
Data_fa$sWM_change<-Data_fa$sWM.y - Data_fa$sWM.x

Data_fa<-Data_fa[which(!is.na(Data_fa$pWM_change)),]

ends<-read_excel('./fa_volume.xlsx',sheet = 'volume')
Data_fa<- left_join(Data_fa,ends,'sub')

Data$time<-factor(Data$time,levels = c("acute","chronic"))
tiff('./sWM_behavior.tiff')
ggplot(data = Data,aes(x=time,y=sWM))+
  geom_boxplot(color='gray',lwd=2,width=0.25)+
  geom_point(size=1,color='black')+
  geom_line(aes(group=sub,color=sub),size=1,linetype=2)+
  theme_bw()+
  theme(legend.position = 'none')+
  theme(axis.title = element_blank())+
  theme(axis.text.x = element_blank())+
  theme(axis.text = element_text(size = 20,face = "bold"))+
  ylim(0,7)
dev.off()

RP<-data.frame(matrix(NA,7,2),row.names = Fa_name[c(1,3,5,9,11,13,15)])
colnames(RP)<-c("r","p")
for (ii in 1:7) {
  jj<-Fa_name[c(1,3,5,9,11,13,15)][ii]
  tmp<-Data_fa[which(!is.na(Data_fa[jj])),]
  Res<-pcor.test(tmp$sWM_change,tmp[jj],tmp[c(3,43+ii)],method = "pearson")
  RP[ii,"r"]<-Res$estimate
  RP[ii,"p"]<-Res$p.value
}
round(RP,3)
colSums(!is.na(Data_fa[21:39]))

RP_L<-RP[c(1,2,3,5,6),]
RP_L$p_fdr<-p.adjust(RP_L$p,'fdr')
RP_L

tmp3<-Data_fa[which(!is.na(Data_fa["ILF_L"])),]
tmp3$sWM<-resid(lm(sWM_change~sWM.x+ILF,tmp3))
tmp3$FA<-resid(lm(ILF_L~sWM.x+ILF,tmp3))
tiff("./corr_ilf_sWM.tiff")
ggscatter(tmp3, x = "FA", y = "sWM",
          add = "reg.line", conf.int = TRUE,
          xlab = 'Left ILF Residual',ylab = 'sWM Residual',
          cor.coef = F,ylim=c(-3,2),xlim=c(-0.065,0.065) )+
  theme(text=element_text(size=20,face = "plain"))
dev.off()
```
